# Individual differences in ethanol drinking meal structure are shaped by social environments

**DOI:** 10.64898/2026.06.17.732974

**Authors:** Marie A. Doyle, Caitlyn M. Edwards, Sabrina D. Hallal, Samuel M. Bond, Nicholas Petersen, Danny G. Winder

## Abstract

Alcohol use disorder (AUD) is marked by substantial heterogeneity in drinking behaviors and health outcomes, underscoring the need for preclinical models that capture interindividual variability. We recently developed open-source capacitive lickometer systems for high-resolution monitoring of mouse fluid intake. Using LIQ PARTI and LIQ HD, we found substantial individual differences in alcohol intake that varied across sex and housing status in C57Bl6/J mice. Here, we conducted a secondary analysis of this continuous access ethanol drinking data to quantify behavioral variability in group and singly housed mice. We introduce a fluid “meal” pattern analysis that integrates drinking across ethanol and water sippers to define discrete drinking episodes. Using this approach, we observed sex- and housing-dependent reorganization of drinking structure across group and single-housed settings, with group-housed male mice exhibiting fewer but faster liquid meals. To further characterize multidimensional drinking patterns, we applied principal component analysis to meal variables and identified a “distributed meal” phenotype defined by increased meal number, reduced meal size, earlier onset of drinking, and higher ethanol preference. Considering factors that influence behaviors in a social environment, we next examined whether social hierarchy was associated with these patterns using a tube test dominance assay. Social rank was unrelated to ethanol and meal measures; however, offensive dominance behavior positively correlated with principal component scores in males. Together, these findings demonstrate that high-resolution, longitudinal analysis of ethanol drinking reveals distinct behavioral phenotypes that are associated with key components of social behaviors, providing a potential framework for understanding heterogeneity in AUD-related drinking.

**Highlights:**

- LIQ PARTI and HD enable high-resolution analysis of ethanol drinking patterns.
- Fluid meal analysis captures sex- and housing-dependent drinking structures.
- Behavioral phenotyping reveals individual differences beyond total ethanol intake.
- PCA identifies a meal phenotype associated with male offensive dominance behavior.

## 1. Introduction

Alcohol Use Disorder (AUD) is marked by substantial heterogeneity in alcohol drinking behavior, spanning total intake levels, drinking patterns, and drinking setting (social versus solitary) [1–3]. In the clinical population, these drinking profiles have been associated with disparate health trajectories and outcomes, such as findings that patient subgroups linked to solitary drinking behavior tend to have worse treatment outcomes [4]. Individual differences in ethanol consumption are also observed in preclinical rodent models [5–15], with a current emphasis on aspects of ethanol drinking and predictors of health outcomes. Given the known heterogeneity in drinking behaviors, mouse models identifying the sources of variability and characterizing social environment influences on drinking enhances translational fidelity to the clinical literature and provides deeper insight into AUD.

In mouse models of AUD, individual variability in ethanol drinking has been associated with a broad set of factors, including strain, age, prior ethanol exposure, stress responsivity, and housing setting. For instance, C57BL/6J mice consume more ethanol than many other strains [16, 17], and high basal anxiety-like behavior is predictive of higher ethanol preference and intake [11]. Additionally, sex consistently accounts for a large share of variance across drinking paradigms, with female C57BL/6J exhibiting higher voluntary intake across a range of drinking protocols relative to males [18–27], a finding which appears to largely generalize to rodent drinking behaviors [17, 28–30]. One variable understudied in preclinical work is social environment. Due to the logistical necessity of singly housing animals in most volitional intake tasks, social aspects of group housing, such as social hierarchy, remain largely uncharacterized in the mouse literature. However, evidence from nonhuman primates and other socially organized rodents, though mixed, largely points towards an inverse relationship between social rank and ethanol intake [31–33], suggesting that social dynamics may play an important role in shaping ethanol intake behavior. Taken together this literature supports further evaluation of sex and social influences in group housed mice to delineate their contributions to individual differences in voluntary ethanol intake.

To facilitate investigation of these factors, our group recently developed a suite of open-source capacitive lickometer tools for high-resolution monitoring of fluid intake. A previous study developed and validated Lick Instance Quantifier with Poly-Animal RFID Tracking Integration (LIQ PARTI) [34], an open-source RFID-based lickometer for group-housed mice, and used it alongside the related single-housed mouse system Lick Instance Quantifier Home Cage Device (LIQ HD) [35] to determine how sex and housing conditions (group vs single housing) influenced ethanol intake and bout microstructure on a population level. Using these novel technologies, we replicated greater ethanol intake in female mice across housing conditions and characterized robust increases in ethanol intake during social isolation in both sexes, along with associated changes in bout microstructure [34]. Although individual differences were not examined in the original study, qualitative observations of variability in ethanol intake motivated further analysis of interindividual variation in drinking patterns and their potential sources.

To address these points, we conducted a secondary analysis of this previously published dataset [34]. Expanding from the original publication’s population-level analysis and a focus on individual lick and drinking bout microstructure findings, we now assess sources of interindividual variability in ethanol consumption as well as add additional drinking pattern analysis. Specifically, we introduce a fluid meal pattern analysis which allows for quantification of drinking episodes across both water and ethanol sippers, finding a sex- and housing-dependent reorganization of meal structure in which group housing resulted in fewer, faster meals in male mice. To further quantify and interpret these multidimensional drinking data, we applied principal component analysis (PCA) to meal data and identified a “distributed meal” drinking phenotype characterized by greater meal number, smaller meal size, earlier initiation after dark onset, and higher ethanol preference. When next considering factors that may influence drinking behaviors in a social, group-housed setting, we chose to focus on social hierarchy. Tube test assays were used to identify social rank within cages, and interestingly, social rank was largely uncoupled from drinking behaviors; however, offensive dominance behavior correlated with drinking PC1 scores in male mice. Further, there was no association between social rank and drinking order during group housed drinking assays. Together, these results demonstrate the value of examining individual differences in ethanol drinking patterns through longitudinal, high-resolution measures across biological and social settings, providing new insight into the heterogeneity of AUD.

## 2. Methods

We conducted a secondary analysis of a previously published mouse continuous access ethanol drinking dataset reported in Petersen et al. [34]. The original study developed and validated LIQ PARTI, an open-source RFID-based lickometer for group-housed mice and used it alongside a single-housed system (LIQ HD) [35] to determine how sex and housing conditions (group vs individual housing) influenced ethanol intake and bout microstructure on a population level.

### 2.1. Animals

In the Petersen et al., 2024 study, male (n=10) and female (n=10) C57BL/6J mice (8 weeks of age) were purchased from Jackson Laboratory (#000664). Mice were housed 5 mice per cage during group housed phases of experimentation and allowed to habituate to the facility for at least 7 days prior to the start of behavioral testing. Mice were housed on a 12 hr light-dark cycle at 22-25°C with *ad libitum* water and food. Behavioral assays took place during the light phase. All experiments were approved by the Vanderbilt University Institutional Animal Care and Use Committee (IACUC) and were carried out in accordance with the guidelines set in the Guide for the Care and Use of Laboratory Animals of the National Institutes of Health.

### 2.2. Overview of continuous access drinking

Detailed methods for continuous access drinking are provided in Petersen et al. [34]. Briefly, continuous access two bottle choice ethanol drinking was measured across a two-staged sequence. In a group housed setting with a LIQ PARTI device and had access to only water for 7 days, 3% ethanol for 3 days, 7% ethanol for 7 days, and 10% ethanol for 4 weeks. Animals were then individually housed with a LIQ HD system and provided access to 10% ethanol for 3 weeks. Bottles and mice were weighed at the end of each 3-7 day recording period with bottle positions swapped to control side bias. Recordings resumed immediately after measurements.

### 2.3. Meal drinking structure analysis

Meals were defined by an inter-response interval threshold of 300 seconds (5 min), such that an individual mouse did not lick at either the ethanol or water bottle for a minimum of 300 seconds. This threshold was selected based on meal definitions set previously [36, 37]. Number of meals, average meal size (licks per meal), meal length (duration of meal), latency to first meal following dark onset, intermeal interval, ingestion rate (meal size/meal length), and ethanol preference within a meal were quantified using data from the dark cycle only. All other drinking calculation and microstructure analyses are calculated as described in Petersen et al. [34].

### 2.4. Tube Test Dominance Assay

In addition to the behavioral assays outlined in Petersen et al. [34], animals underwent tube test dominance testing as follows, adapted from Harrison et al. [38, 39]. Prior to the start of dominance testing, animals were habituated to the tube apparatus by placing the animal at one end and allowing them to escape out the other end three times. All animals readily acquired this task. During dominance testing, mice were tested pairwise in the tube and the trial was ended when one mouse retreated and all 4 paws were outside of the tube. For all mice this took less than 3 minutes, so no trials were excluded from analysis. Pairings were conducted in a round-robin design to randomize test order, with at least 3 minutes of rest in the home cage between a mouse ending one trial and starting their next. Three round-robin series were conducted per day, with at least 1 hour of rest between each. Animals were only paired with cage mates. Trials were recorded using an overhead camera for behavioral analysis.

Behaviors were hand-scored frame-by-frame using the open-source software BORIS [40]. For each contest, the start was defined as the moment when the experimenter released both mice into the tube, and the end was defined as the moment when one of the two mice had all four paws outside the tube, as previously described [39]. The elapsed time between contest start and end was calculated as the contest’s tube time. Four main behavioral states were quantified for each contest: Pushing, Resisting, Forced Retreating, and Voluntary Retreating [39, 41, 42]. Pushing, an offensive move, was defined as a behavior in which the mouse thrusts its head forward at the opponent. Each offensive push was countered by a defensive behavior of resist or forced retreat. Mice were defined as resisting when they pushed back at the offensive mouse and/or held their territory. Forced retreats were identified as events in which the defensive mouse retreats backwards in response to an opponent’s push. Lastly, voluntary retreats were defined as retreating behaviors without an opponent pushing. As contests vary in their total tube time, each mouse’s time spent performing each behavior (pushing, resisting, forced retreating, and voluntary retreating) was quantified as a percentage of their total tube time (e.g., time spent pushing (in seconds) * 100 / total tube time). Cage ranks were determined by calculating each mouse’s cumulative wins across all contests. The mouse with the most wins within the cage was ranked #1, second-most wins received rank #2, etc. In the event of a tie in cage rank, the two mice’s head-to-head win/loss record was determined, and the mouse with more head-to-head wins was given the more dominant rank.

### 2.5. Statistics

#### 2.5.1. Principal Component Analysis

Principal component analysis (PCA) was performed to better understand relationships between drinking variables within our dataset, adapted from previous methods [24, 43–45]. Meal data was binned so that each data point is representative of one subject during on experimental phase, and for this analysis two experimental phases were chosen, the final week (week 4) of group housed drinking at 10% ethanol and the final week (week 3) of social isolation at 10% ethanol. A total of five meal variables were selected using correlational analysis and data from both males and females was combined, resulting in a 20 (subjects) x 5 (variables) matrix to input into the PCA for each experimental week. Principal components were deemed significant based on the Kaiser rule. K-means clustering was performed on the resulting principal component scores; the analysis was run for 2-10 clusters, and the final number of clusters was chosen based on silhouette scores. PCA and K-means clustering were performed with custom Python code in a Jupyter Notebook.

#### 2.5.2. Markov Models

To identify any effect dominance may have on ethanol drinking order within each cage, we utilized Markov models to calculate the probability of transitioning from one state to another, with transitioning to a given state solely dependent on the previous state [46]. These probabilities are represented using transition matrices to describe the probability of one mouse drinking after another within 15 seconds for bouts and 400 seconds for meals to show order of drinking events. These transitions were determined for dark cycle drinking during both the water week and for week 4 of group housed drinking at 10% ethanol. For both time points, dominance ranks from the tube test assays were used to indicate dominance. Transition matrices were calculated using Python code in a Jupyter Notebook, using the *mchmm* package.

#### 2.5.3. Other statistical analyses

Full statistical analyses and results are listed in Supplemental Table 1. Statistical analyses were performed using GraphPad Prism or custom Python code in a Jupyter Notebook, and all values are represented as mean ± SEM, unless otherwise noted. An unpaired t-test (two-tailed) was used to compare the means of two groups and a one-way analysis of variance (ANOVA) was used to compare the means of three groups, followed by a Tukey post-hoc test when appropriate. A two-way ANOVA with repeating measures (RM) was used when comparing two independent variables with repeated measures, followed by a Sidak post-hoc test when appropriate. Post hoc statistical hypothesis testing accounted for multiple comparisons in appropriate cases. Correlation analyses were specified *a priori*, where Pearson correlations and linear regressions assessed relationships and multiple comparisons were accounted for using a False Discovery Rate (Benjamini and Hochberg). Significance was defined as *p<0.05 and **p<0.01, unless otherwise noted.

## 3. Results

### 3.1. LIQ PARTI captures stable inter-individual differences in ethanol preference and circadian drinking patterns

In the original Lick Instance Quantifier with Poly-Animal RFID Tracking Integration (LIQ PARTI) publication, Petersen et al. analyzed ethanol drinking patterns in the form of individual licks (number, frequency, duration, and inter-lick intervals) and drinking bouts (number, size, duration), with comparisons made on a population level [34]. Briefly, animals were first group housed with a LIQ PARTI device and had access to 10% ethanol for 4 weeks following a week of water-only drinking and an ethanol ramp. Mice were then individually housed with a Lick Instance Quantifier Home Cage Device (LIQ HD) system and provided access to 10% ethanol for 3 weeks (Figure 1A). New visualization of individual ethanol preference and its trajectory across drinking stages revealed substantial inter-individual variability (Figure 1B), with significant variability observed across ethanol dose, housing condition, and sex (Levene’s test, Dose: 4.82, p=0.004; Housing: 8.60, p=0.0056; Sex: 14.97, p=0.0002). Ethanol preference was not random and instead reflected stable individual drinking patterns, as average ethanol preference during the first week of 10% ethanol access was significantly correlated with preference during the final week in both group housed (Supplemental Figure 1A, R^2^=0.74, p<0.001) and socially isolated (Supplemental Figure 1B, R^2^=0.57, p<0.001) settings, respectively. These observations motivated further analysis of drinking patterns and their potential sources.

**Figure 1.**
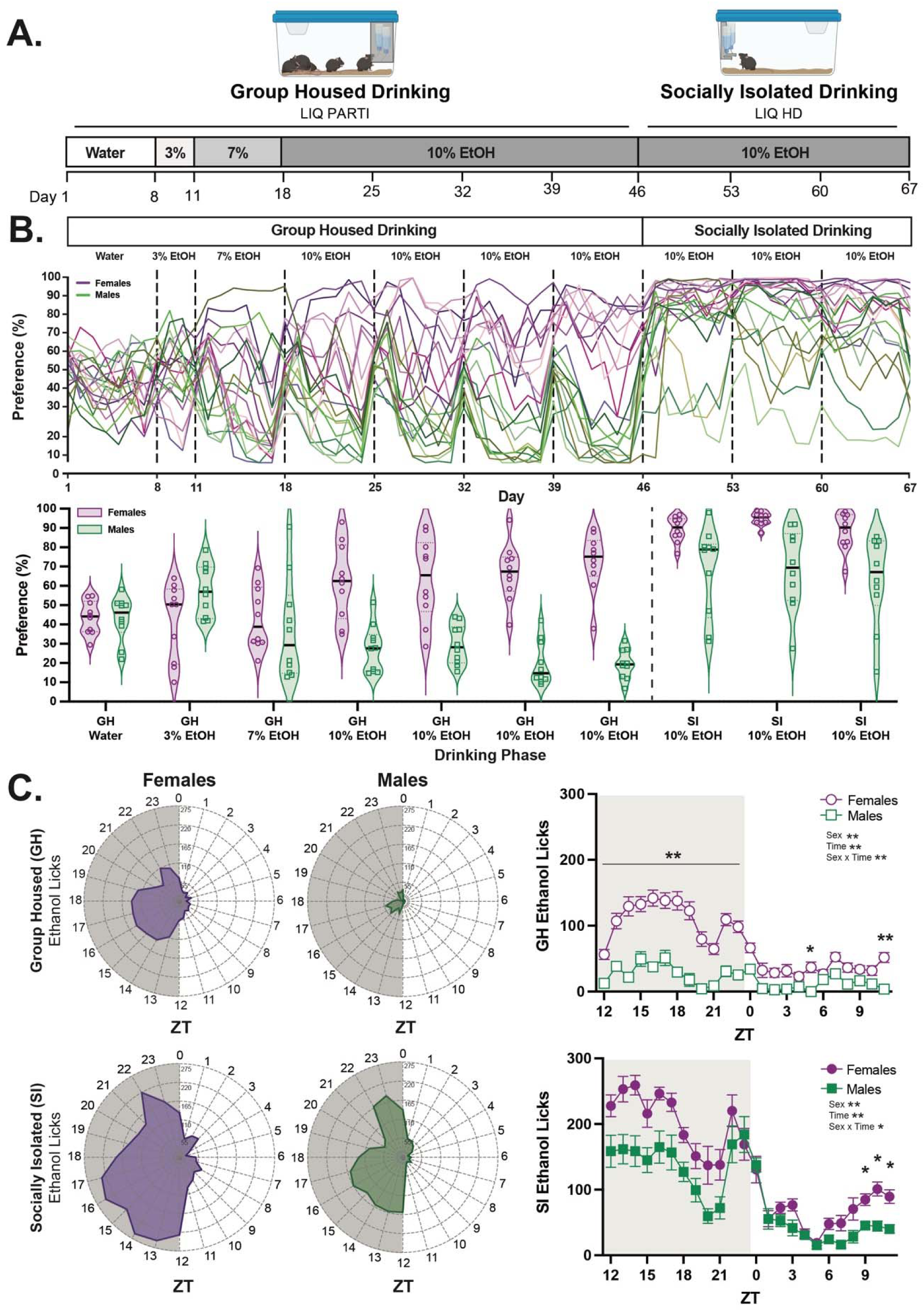
LIQ PARTI and LIQ HD systems capture heterogeneity in ethanol preference and circadian drinking patterns. **A.** Experimental timeline for continuous access ethanol drinking. **B. Top:** Representation of interindividual variability of ethanol preference calculated from total lick number at each sipper, where each line represents an individual animal (n=10/sex); **Bottom:** Significant variability was observed across ethanol dose, housing condition, and sex in average ethanol preference calculated from total lick number at each sipper (n=10/sex, Levene’s test). **C. Left:** Chronograms of average daily licks at the ethanol bottle across zeitgeber time (ZT) during the last week of 10% ethanol access during group housed (top) and socially isolated (bottom) drinking (n=10/sex); **Right:** Sex differences in circadian patterns of ethanol licks were observed across zeitgeber time (ZT) during the last week of 10% ethanol access during group housed (top) and socially isolated (bottom) drinking settings (n=10/sex, two-way ANOVA with repeated measures). All data are represented as mean ± SEM, with significance was defined as *p<0.05 and **p<0.01.

Petersen et al. previously reported that female mice consume more ethanol than males in both group housed and socially isolated settings but focused on daily or weekly consumption measures [34]. Here, data were first examined as licks at the ethanol bottle across a 24 hour cycle to assess circadian patterns of intake (Figure 1C). In a group housed setting, females licked at the ethanol bottle more than males across the entirety of the dark (active) cycle as well as at the very end of the light (inactive) cycle (Figure 1C). When singly housed, both sexes increased their ethanol intake [34], reducing circadian-related sex differences in intake. Specifically, in contrast to group housed drinking, socially isolated females licked at the ethanol bottle significantly more than socially isolated males only in the three hours preceding dark onset (Figure 1C). Together, these data highlight sex differences in circadian drinking patterns observed in continuous access ethanol intake, particularly those at the end of the light cycle, and support further investigation of drinking structure during dark cycle drinking, consistent with the active phase in mice.

### 3.2. Meal pattern analysis reveals sex- and social setting-dependent organization of ethanol intake

Given the differences in drinking patterns observed across the circadian cycle, we added additional drinking structure analysis in the form of a fluid meal pattern (“meal”) analysis for dark cycle data. These data take a higher-level analysis from bout data to include drinking at both ethanol and water bottles grouped into a single drinking episode, where meals were defined as drinking at either sipper within an inter-response interval threshold of 300 seconds [36, 37] (Figure 2A). Broadly, we found that sex and housing condition both contributed to changes in meal structure. First, we focused on meal number metrics. While we observed no sex differences in cumulative meal number across the dark cycle during the water-only week (Figure 2B), with the addition of ethanol availability, female mice displayed significantly greater cumulative meals compared to males in both group housed and socially isolated settings (Figure 2B). These differences were driven by decreased meal number in males during group housing in contrast to an increase in meal number in females during the social isolation drinking, both compared to their respective water-only meal number (Figure 2C). These findings suggest a sex- and housing-dependent reorganization of meal patterns, with males and females exhibiting distinct adaptations to ethanol availability.

**Figure 2.**
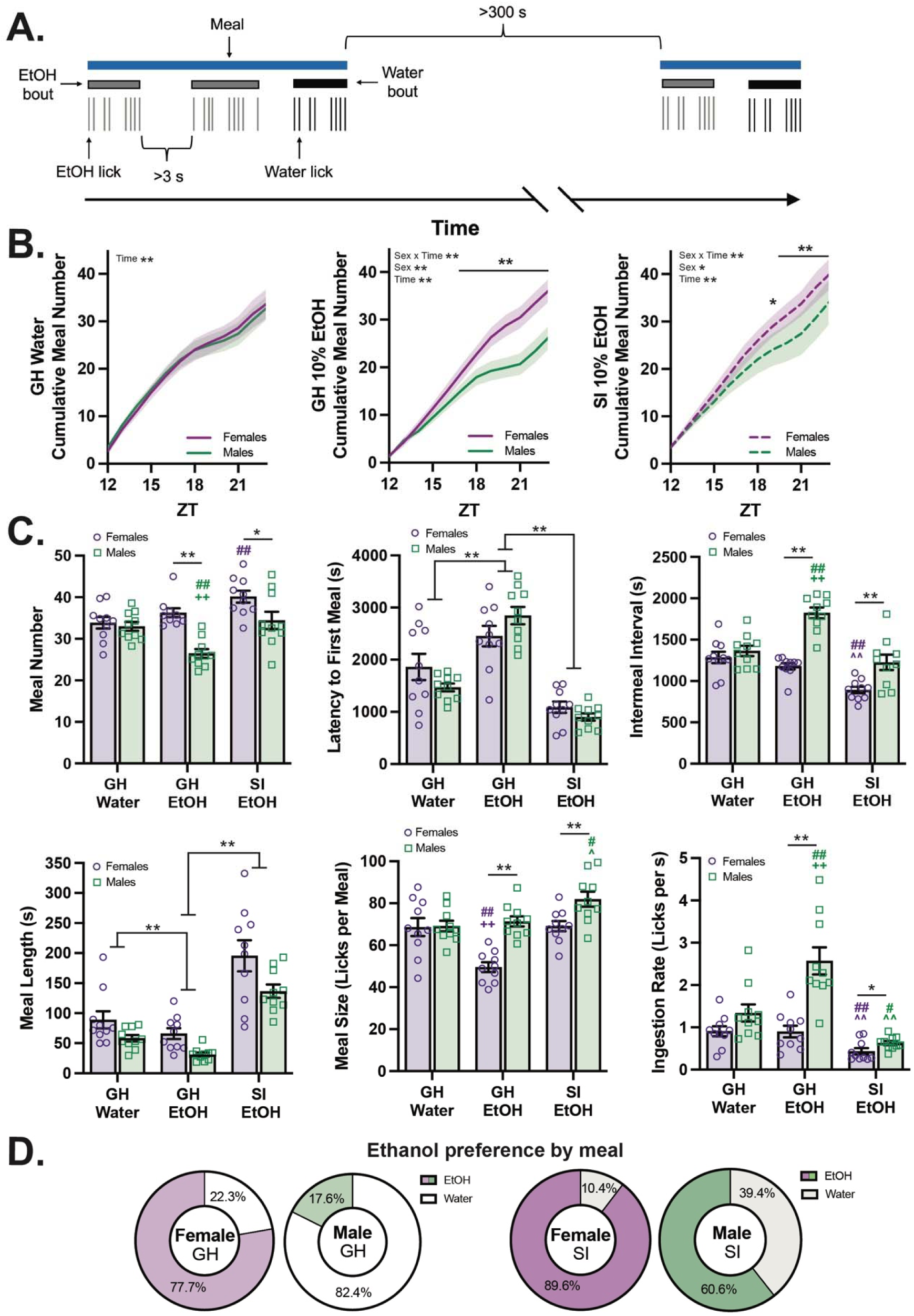
Meal pattern analysis identifies sex- and social environment-dependent features of ethanol intake. **A.** Visual representation of a fluid meal in relation to lick and bout events. **B.** Female mice displayed significantly greater cumulative meals across the dark cycle compared to males in both group housed and socially isolated settings (n=10/sex, two-way ANOVA with repeated measures). Data are represented as mean ± 95% confidence interval. **C.** Analysis of meal variables meal number (main effects of time and sex, interaction of time x sex), latency to the first meal in the dark cycle (main effect of time), intermeal interval (main effects of time and sex, interaction of time x sex), meal length (main effects of time and sex), meal size (main effects of time and sex, interaction of time x sex), and ingestion rate (main effects of time and sex, interaction of time x sex) revelated robust sex- and housing setting-dependent aspects of ethanol drinking. (n=10/sex, two-way ANOVA with repeated measures). * (black) represents a significant difference between sexes at the labeled timepoint when there was a significant interaction of sex x time. **#** represents a significant difference from the water timepoint of the corresponding sex (purple for females, green for males). **^** represents a significant difference from the group housed ethanol timepoint of the corresponding sex (purple for females, green for males). **+** represents a significant difference from the socially isolated timepoint of the corresponding sex (purple for females, green for males). **D.** Ethanol preference calculated from meal drinking data for male and female mice in both group housed and socially isolated drinking settings (n=10/sex). Data are represented as mean ± SEM with significance was defined as *p<0.05 and **p<0.01, unless otherwise noted.

We next extended meal structure analysis to include latency to the first meal in the dark cycle, intermeal interval, meal length (duration of meal), meal size (number of licks per meal), and ingestion rate (meal size/meal length). While latency to first meal was not different between sexes, there was a significant main effect of time. Here, there was a significant increase in latency during group housed ethanol drinking followed by a significant decrease in latency during social isolation ethanol drinking, both compared to the water-only week (Figure 2C). Intermeal interval showed the inverse pattern to meal number such that males show an increased intermeal interval during group housing with ethanol whereas females show a decreased intermeal interval during social isolation. Sex differences in meal number and intermeal interval were reflected in meal length and size metrics. Here, a main effect of time showed decreased meal length during group housed ethanol drinking and increased meal length during socially isolated meal drinking (Figure 2C). A main effect of sex displayed longer meals overall in females compared to males (Figure 2C). Additionally for meal size, female mice decreased their meal size during group housed drinking while males increased meal size during social isolation, where sexes are significantly different from each other at each timepoint (Figure 2C). Taking meal length and size together to estimate the average ingestion rate during a meal, we found that group housed males during ethanol greatly increase their ingestion rate compared to their water week and compared to females during the same time point. We also find that both male and female animals drink more slowly in a meal once in social isolation. Finally, meal data also reflected lick- and bout-level data for calculating ethanol preference (Figure 2D), as these numbers were consistent with previously reported ethanol lick and bout values [34]. Taken together, meal data allow for a more integrative assessment of intake structure across bottles, enabling a more complete perspective on drinking behavior, and further detection of sex- and housing-dependent shifts in drinking organization that were not apparent from lick- or bout-level analyses alone. Females show consistent, robust ethanol drinking and preference across social setting, whereas males appear highly sensitive to social environment.

### 3.3. A sex-defined “distributed meal” phenotype emerges from multivariate drinking structure

Given the sex- and social setting-dependent reorganization of meal structure and ethanol preference, we next used a multivariate framework to explore meal factors that govern the substantial observed individual differences in ethanol behaviors (Figure 1B).To quantify and interpret these multidimensional behavioral data, we applied principal component analysis (PCA) to drinking meal data obtained during the final week of 10% ethanol drinking in both the group-housed and socially isolated phases. PCA allowed us to reduce meal structure and preference variables into orthogonal components that capture the primary axes of behavioral variance. Meal data variables reflecting drinking initiation (meal number and latency following dark onset) and termination/satiety (meal size and meal length) as well as preference for ethanol within the meal were included to identify coordinated patterns of drinking organization rather than isolated metrics (Figure 3A). Intermeal interval and ingestion rate were not included due to a high correlation with meal number and dependency on other included variables, respectively.

**Figure 3.**
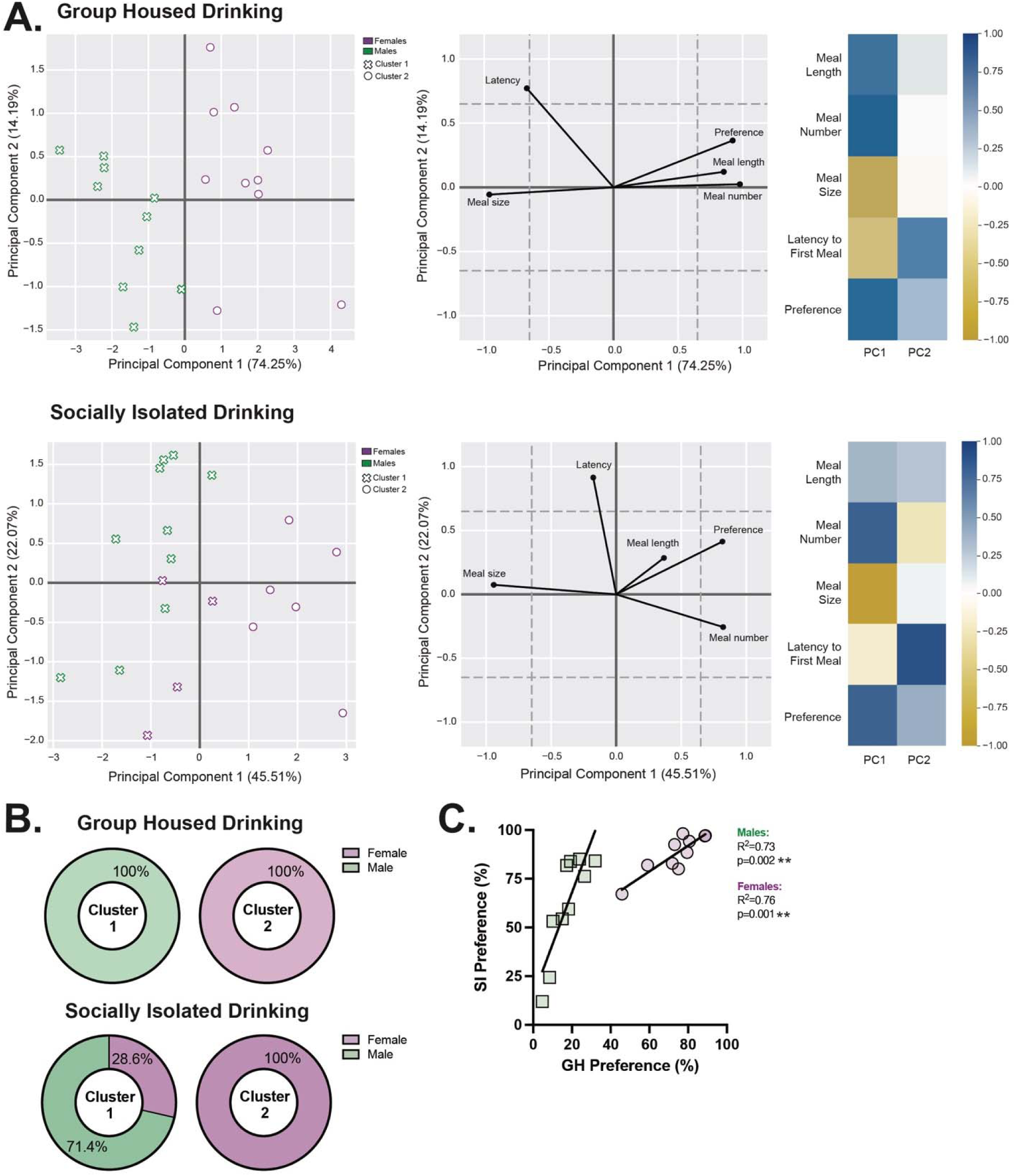
Multivariate analysis of drinking structure reveals a sex-associated “distributed meal” phenotype. **A.** Principal component analysis of meal variables. **Left:** Score plot of individual animals; **Center:** Loading plot of meal variables; **Right:** Loading heatmap of meal variables. **Top:** In group housed drinking, ethanol preference, meal length, and meal number loaded positively and meal size and latency loaded negatively on PC1; **Bottom:** In socially isolated drinking, ethanol preference and meal number loaded positively and meal size loaded negatively on PC1, and latency loaded significantly onto PC2 (n=10/sex). **B.** K-means clustering defined two clusters for each housing condition’s PCA, which was primarily defined by sex. **C.** Ethanol preference calculated from meal analysis data was significantly correlated between the two housing conditions in both male and female mice (n=10/sex, simple linear regression). All data are represented as mean ± SEM, with significance was defined as *p<0.05 and **p<0.01.

Across both housing conditions, variables loaded onto principal components (PCs) 1 and 2 in similar patterns (Figure 3A). Loadings in group housing were particularly strong, where ethanol preference, meal length, and meal number loaded positively and meal size and latency loaded negatively on PC1. During social isolation, ethanol preference and meal number again loaded positively and meal size loaded negatively on PC1, but meal length did not strongly load onto PC1. Latency instead loaded significantly onto PC2 only. We then applied k-means clustering to the PCA scores to determine whether animals segregated into distinct behavioral groupings (Figure 3A). The optimal solution identified two clusters, in which both group housed (Fisher’s exact test: p<0.01) and social isolation (Fisher’s exact test: p=0.011) settings were defined by sex (Figure 3B). Finally, though meal structures changed across housing conditions, ethanol preference between the two was highly correlated, again suggesting stable individual drinking trajectories (Figure 3C). In combination with data from Figure 2, these findings indicate social setting-associated shifts in drinking behavior, where group housing with ethanol access suppresses male meal initiation and continuity resulting in initiation of fewer, faster meals. We also observed a shift in meal length such that both sexes take longer meals when socially isolated compared to group housed drinking. Together, the dimensionality reduction with PCA further revealed a “distributed meal” behavioral phenotype, reflecting coordinated organization of drinking characterized by ethanol preference, more meals across the dark phase, smaller meal size, and earlier initiation after dark onset. These features describe a distributed, frequent-initiator pattern of ethanol-preferring intake rather than consolidation into a few large episodes in which sex is a primary driver of individual variability.

### 3.4. Meal phenotypes are associated with key aspects of dominance behavior

One possible interpretation of the observed drinking behavior is that group social environments induce competition that shift meal initiation and structure based on dominance, hypothetically with reduced sipper access encouraging short, rapid meals. To examine this, we characterized the relationship between the principal component scores with social hierarchy and dominance behaviors. Given evidence from the primate literature linking ethanol intake to social rank [32, 33], we examined whether dominance status related to drinking behaviors in our cohort. Social hierarchy within each cage was assessed using the tube test dominance assay, in which cage mates were placed at opposite ends of a narrow tube and the trial concluded when one mouse retreated fully, designating the remaining animal as the winner of the bout. Dominance testing was conducted following completion of the group housed drinking phase (Figure 4A), with social rank defined by the highest percentage of wins (Figure 4B). Interestingly, there were no differences in ethanol preference across social rank (Figure 4C). In addition to overall wins, we further analyzed tube test behaviors including percent of time spent pushing, resisting, voluntarily retreating, and being forced to retreat. Though these metrics have been previously correlated with social rank in the tube test, no significant differences across rank were observed in our data set (Figure 4B). We then asked whether individual values for these measures of social dominance related to multivariate drinking behavior by correlating them with PC1 scores from the group housed and socially isolated drinking analysis. No significant associations were observed between the “distributed meal” phenotype and percent win (Figure 4D). When other dominance measures were correlated with PC1 scores, there was a significant correlation between the one offensive behavior measured, push time, in male but not female mice (Figure 4D). This correlation was not present with socially isolated PC1 scores (Figure 4D). No other significant associations were found with the defensive behaviors (Supplemental Figure 2), suggesting that heterogeneity in meal structure occur largely under offensive dominance behavior initiation. Together, these findings suggest that individual differences in meal phenotypes are influenced by social interactions, with offensive dominance behavior, but not tube test-derived social rank or defensive behaviors, associated with drinking phenotypes in males.

**Figure 4.**
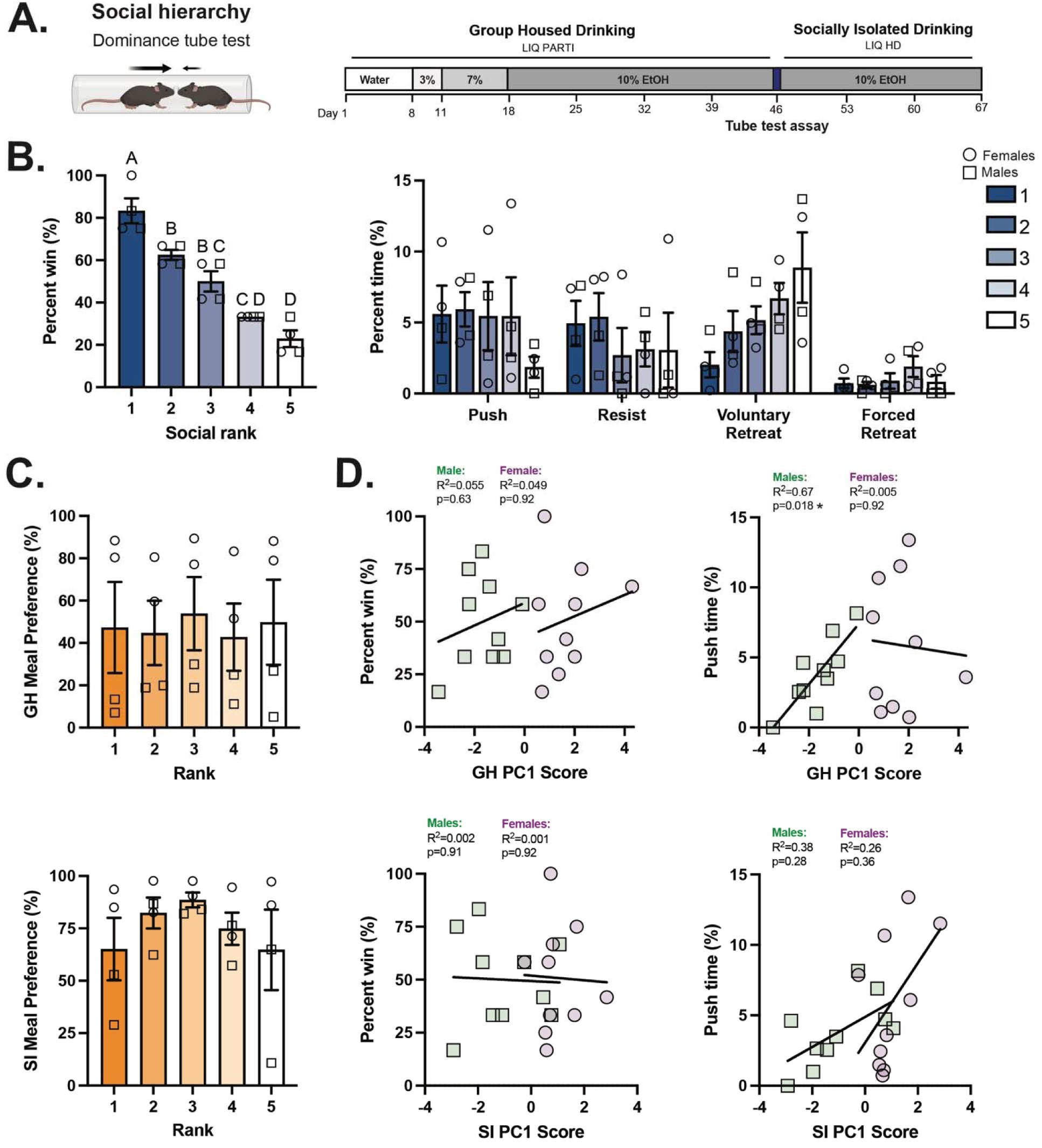
Meal phenotypes are associated with specific features of dominance behavior but not social rank. **A.** Visual representation of tube test dominance assay and timeline of testing across the continuous access drinking paradigm. **B. Left:** Social rank is defined by percent win during the tube test dominance assay; **Right:** Additional variables of time spent pushing, resisting, voluntarily retreating, and being forced to retreat were not significantly different across social rank (n=10/sex; n=4 cages, One-way ANOVA). **C.** No significant differences in ethanol preference were observed across social rank in either group housed or socially isolated drinking measures (n=10/sex; n=4 cages, One-way ANOVA). **D. Left:** Percent win, the metric for defining social rank, was not significantly correlated with PCA-defined drinking phenotypes (PC1 scores) in either housing environment; **Right:** The offensive dominance behavior of pushing was significantly correlated with PCA-defined drinking phenotypes (PC1 scores) only in group housed males (n=10/sex, Simple linear regression with FDR correction). All data are represented as mean ± SEM, with significance was defined as *p<0.05 and **p<0.01.

### 3.5. Social hierarchy status did not govern drinking order

Beyond overall ethanol intake and drinking patterns, we also asked whether the order in which mice accessed the drinking bottles might reflect social hierarchy. While metrics related to meal patterning could be independent of social rank, the sequence of drinking could still reveal subtle dominance-related behaviors. To explore this, we applied Markov modeling, which provides a framework for analyzing the probability of transitions between discrete events, in this case successive drinking events between cage mates, allowing us to capture patterns in drinking order analyzed during the final week of group housed 10% ethanol access. We initially applied this approach to meal-level data; however, these longer recording periods likely obscured effects (Supplemental Figure 3). We therefore focused on drinking bouts. Given the strong and opposite preferences formed by males and females during group housed drinking, we analyzed bouts at the preferred bottle, ethanol for females and water for males. Identical analyses were performed for drinking bouts during the water-only week. Using this approach, we found no consistent associations between social rank and drinking order in either sex (Figure 5), suggesting that dominance does not govern the sequence of drinking behavior during either water or in ethanol drinking sessions. Consistent with the PCA-defined phenotype, the lack of rank-related ordering indicates that the distributed, ethanol-preferring meal phenotype is not implemented through priority in drinking order. Instead, phenotypic differences arise from how individuals initiate and structure bouts within a drinking session independent of dominance-based turn taking.

**Figure 5.**
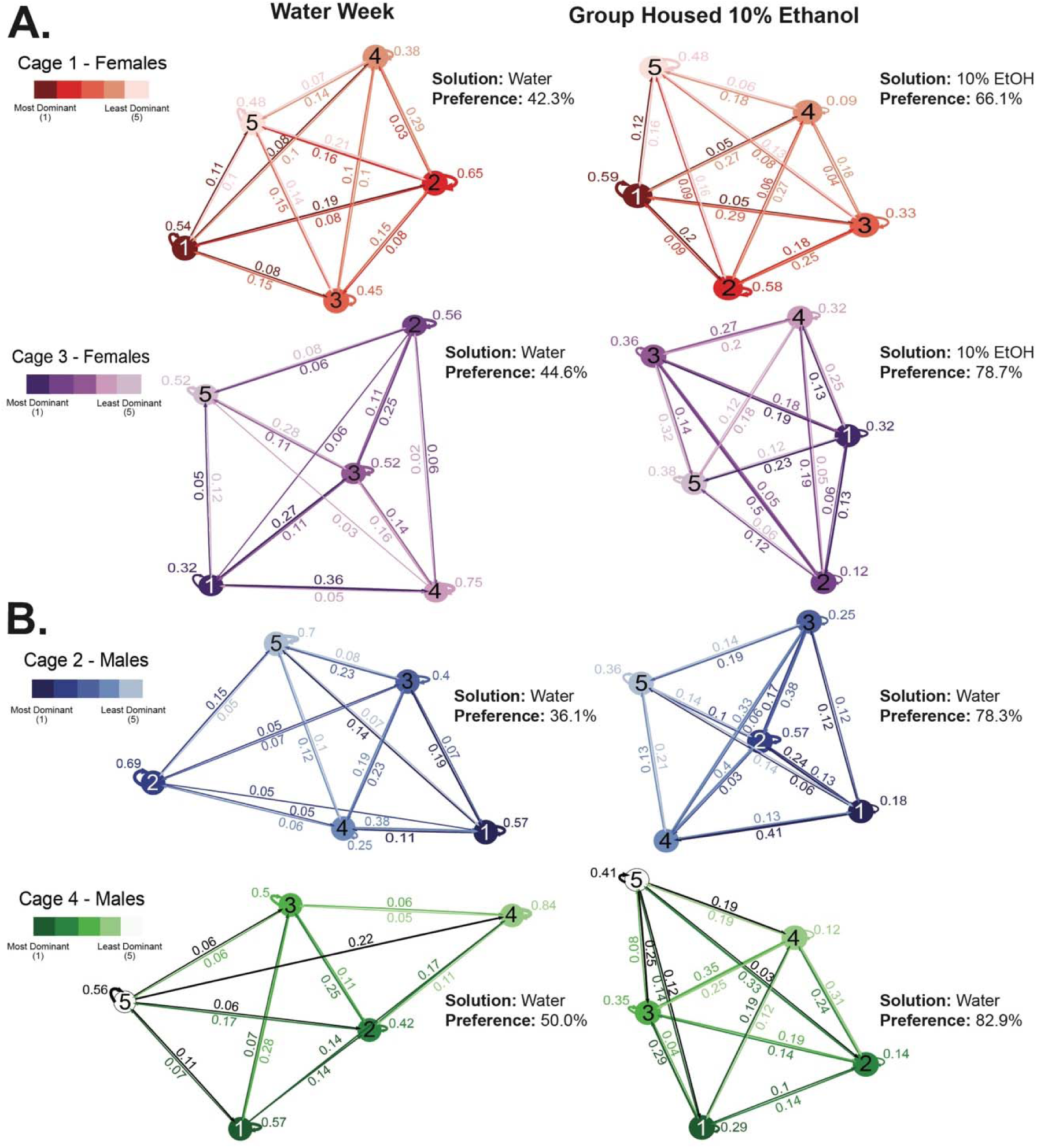
Bout drinking order at the preferred sipper was not predicted by social rank. **A.** Markov modeling of bout drinking order in female mice did not display a relationship between social rank and drinking order. **Left:** Water week; **Right:** Last week of group housed 10% ethanol access (n=5 mice/cage; n=2 cages/sex). **B.** In male mice, Markov analysis of bout drinking order showed no association between social rank and drinking order. **Left:** Water week; **Right:** Last week of group housed 10% ethanol access (n=5 mice/cage; n=2 cages/sex).

## 4. Discussion

Taken together, our results demonstrate profound, stable individual variation present in continuous ethanol access home cage drinking behavior among male and female mice (Figure 1), with fluid meal patterns defining a valuable new level of analysis to identify key aspects of heterogeneity in drinking behaviors. Using meal-level analysis, we first found a sex- and housing-dependent reorganization of meal structure in which ethanol availability in group housing resulted in fewer, faster meals in male mice and social isolation produced an increase in meal number, length, and size in both sexes (Figure 2). Next, dimensionality reduction with PCA further revealed a “distributed meal” behavioral phenotype in which sex is the main driver of individual variability in drinking behavior across the experiment (Figure 3A-B). Given this interpretation of drinking phenotype and how social aspects of drinking are a contributing factor to AUD pathogenesis in the clinical population, we tested how within-cage social hierarchy affects drinking behavior. We observed a significant association between one proactive dominance-related behavior, push time, and “distributed meal” drinking behavioral phenotype (PC1 scores), suggesting that mice displaying greater offensive dominance behavior exhibit a more relaxed, sustained drinking strategy and do not need to constrain their drinking based on the social environment (Figure 4D). Finally, Markov modeling demonstrated that social rank has limited effect on specific drinking order among group-housed mice (Figure 5). These findings establish fluid meal analysis as a powerful approach for resolving individual differences in ethanol drinking behavior, identifying sex as the primary source of variability and highlighting social setting as a critical axis for future mechanistic dissection and translational modeling of alcohol-related behaviors.

### 4.1. LIQ PARTI models heterogeneity in ethanol drinking behavior

Our study has demonstrated that the LIQ PARTI and LIQ HD home cage lickometer systems [34, 35] can reveal clinically-relevant heterogeneity in drinking patterns, as demonstrated through the substantial interindividual variability in ethanol preference (Figure 1B) and meal drinking patterns (Figures 2–3). Individual differences in drinking behavior among AUD patients are well-described but remain understudied, highlighted by the lack of broad-based efficacy among current approved AUD pharmacotherapies. Clinicians and researchers alike have long recognized that heterogeneity exists among AUD patients [2, 3, 47–49], which can be attributed to factors such as drinking patterns, genetics, sociological/cultural factors, and co-morbidities [1, 3, 47–50]. For example, recent work suggests that AUD patients fall into three distinct subgroups based on their anxiety and depression trajectories during early abstinence [1]. At the drinking pattern level, key effectors of AUD heterogeneity include the consistency of heavy or binge drinking, sensitivity to alcohol’s rewarding or aversive properties, and the age of drinking onset [50, 51], critical as distinct drinking patterns are associated with overall health outcomes [52, 53]. Heterogeneity in drinking behavior is also reflected in preclinical animal models [5–8, 12, 13, 54], with a current emphasis on aspects of ethanol drinking and predictors of future health outcomes. To further build on preclinical drinking data, future studies with LIQ PARTI and LIQ HD should evaluate health outcomes (cardiovascular, hepatic, neurologic, etc.) and tissue histology associated with individual drinking patterns to identify subgroup-specific outcomes to inform the development of personalized therapies.

### 4.2. Meals define drinking episodes on a biologically relevant timescale

To examine individual drinking patterns more deeply, we applied meal pattern analysis, a framework widely used in ingestive behavior research, particularly for analysis of feeding [55–58]. In this context, a defining feature of meals is their intentional timing structure, set to encompass satiation and satiety signaling and further distinguishing meals from individual licks or bouts. Specifically, the initiation and conclusion of a feeding meal are governed by satiation and satiety signals, which require time to be communicated to the brain and used to modulate ingestive behavior [59–63]. In the context of ethanol intake, a meal provides a concrete definition of a drinking episode on a timescale that theoretically permits pharmacological feedback from ethanol to influence termination and subsequent initiation of drinking [64, 65]. Focusing on female mice, as most of the drinking occurred at the ethanol bottle in this sex, intermeal interval averaged 16-20 minutes (Figure 2C) compared to interbout intervals of about 1-3 minutes (data not shown). Accordingly, we hypothesize that ethanol consumed per meal would relate more closely to blood ethanol concentration (BEC) measurements than individual licks, bouts, or time-averaged metrics. Specifically, an individual bout would presumably result in much lower BECs that decay faster, failing to represent a true intake epoch. These concepts support a future direction for using meals to interpret the physiological consequences of ethanol intake patterns, with use in determining how meal structure changes during the escalation of ethanol intake to yield important information about the development of excessive drinking, as another example.

### 4.3. Meal pattern analysis reveals shifts in drinking structure by social environment

Meal pattern analysis revealed that females show consistent, robust ethanol drinking and preference across social settings, whereas males appear highly sensitive to social environment (Figures 2C). Specifically, males exhibited fewer, shorter, and faster meals when ethanol was available under group-housed conditions, suggesting that social setting suppresses meal initiation and continuity. A similar profile of fewer, shorter, faster meals has been reported for food intake in group-housed male rats and pigs [66, 67]. A limitation for interpretation is the use of different lickometer systems in the two housing conditions (LIQ PARTI vs LIQ HD). While no known physical constraints in access or acquisition of drinking from different apparatuses exist, it cannot be ruled out that the physical layout of the lickometer systems may contribute to differences in drinking structure.

### 4.4. PCA identifies an “distributed meal” drinking structure with sex as a major driver of variability in drinking

PCA revealed a “distributed meal” phenotype in which higher ethanol preference covaried with greater meal number across the dark cycle, smaller meal size, and earlier initiation after dark onset. Together, these features describe a distributed, frequent-initiation pattern of ethanol-preferring intake. When K-means clustering was applied to PC scores, it separated animals entirely by sex in group housing and largely by sex in social isolation, implying robust, sex-specific drinking strategies embedded in the multivariate space (Figure 3B). This is consistent with the observed univariate differences (Figures 1–2) and highlights that sex remains a principal organizing factor. These sex-dependent results add to the large body of literature indicating that adult female mice drink more ethanol than males across a variety of drinking tasks including continuous access [17, 18, 21–23], drinking in the dark [19, 68], intermittent access [20, 25, 26], and some operant self-administration procedures [21, 69]. The primary driver of higher ethanol consumption among female rodents is still unclear, but some studies indicate a role for sex hormones [70], as ovariectomy significantly reduces female ethanol intake in drinking in the dark and continuous access tasks [71–73]. In addition to circulating hormones, sex chromosomes also play a significant role in sex differences in drinking behavior, with XX mice drinking more than XY, independent of gonad type [74]. However, male mice may be more susceptible to developing greater ethanol intake or aversion-resistant intake following a stressor [8, 75–77], and intriguing area of future investigation. Overall, while prior work has primarily focused on sex differences in total intake and ethanol preference, meal analysis demonstrates the presence of sex-specific drinking patterns as well.

### 4.5. Drinking phenotypes are related to key dominance behaviors specifically in group housed male mice

Due to the logistical challenges of group housed home cage drinking in rodents, few studies have investigated how social hierarchy impacts drinking behavior in mice. Using a tube test dominance assay, we identified within-cage social hierarchies (Figure 4B). Interestingly, PC1 meal structure data positively correlated with percent of time spent pushing in the tube test, a measure of proactive dominance behavior, in group housed males (Figure 3D) but not other metrics, during social isolation, or in females (Figure 3C-D and Supplemental Figure 1). This selective relationship suggests that a specific offensive dominance-related behavioral facet may track with the constellation of “distributed meal” drinking features that define PC1, rather than with general dominance rank. It is possible that these effects in males reflect competition at the sipper during group housing, making it more difficult for individual mice to initiate or sustain a meal. For example, mice may be displaced from the sipper or unable to readily resume drinking, resulting in shorter and smaller meals. However, the absence of predictive relationships in the Markov modeling analysis (Figure 5) suggests that, if access limitations contribute to these effects, they are driven by moment-to-moment competition rather than stable social hierarchy.

Social interactions have a strong and well-known impact on alcohol drinking in the human population [78]. In preclinical models, social aspects of ethanol drinking are often looked at in the context of social hierarchy and dominance. We did not observe a relationship between social rank and ethanol preference in our data set (Figure 4C-D). These results conflict with some studies in non-human primates (NHP) and other rodents [31–33]. In multiple NHP species and in prairie voles, a negative correlation exist between social rank and alcohol intake, whereby subordinate animals drink more ethanol than dominant ones [32, 33], though results are mixed. During a different drinking paradigm, social rank had no impact on ethanol intake among cynomolgus macaques [79]. Interestingly, in rhesus macaques, social aggression is positively correlated with ethanol consumption [80], more closely aligning with our observations and consistent with dominant rats displaying greater efficiency at displacing a cage mate during competition for a resource [81]. It is important to note that drinking sessions in NHP studies were conducted in separate cages; therefore, competition during the session could not contribute to drinking phenotypes.

Moreover, the discrepancy between our findings in mice and some NHP reports may stem from how social dominance was assessed. Dominance measurements differed substantially between preclinical models and are often calculated as a “dominance index,” which incorporates about a dozen ethologically-relevant behaviors into a continuous variable [33]. Our study measured social hierarchy through a round-robin tube test competition, and while widely used [31, 41, 42, 82], it is limited in scope. A more holistic assessment of social dominance [39], incorporating additional dominance assays such as urine marking or warm-spot competition [83], as well as video monitoring and machine learning-based behavioral tracking of mouse-mouse interactions (eg, aggression, allogrooming, competition for sipper access), would provide a more comprehensive representation of social behavior in group-housed mice. Larger cohorts would provide greater statistical power for dominance and social assays as well as support the inclusion of additional variables in future principal component analyses, limitations for the current work. Overall, we propose that social environments induce competition at the sipper that shifts meal initiation and structure, with male mice showing greater sensitivity to this.

## 5. Conclusions

In summary, in this secondary analysis we used a suite of open-source capacitive lickometer tools, LIQ PARTI and LIQ HD, to characterize individual variability in aspects of mouse ethanol drinking behavior and assess the contribution of social environments. A fluid “meal” pattern framework and principal component analysis-based phenotyping further identified distinct individual drinking strategies, primarily associated with sex but not social rank. Future studies are proposed to investigate links between drinking behaviors and health outcome trajectories as well as to better assess social hierarchy in the context of ethanol drinking through holistic approaches and video monitoring. These findings demonstrate that high-resolution analysis of continuous access ethanol intake reveals structured individual variability, providing a framework for studies linking drinking phenotypes to alcohol-related outcomes and the future development of personalized therapeutic targets in AUD.

## CRediT Statement

**MAD:** Conceptualization, formal analysis, funding acquisition, investigation, and writing – original draft preparation; **CME:** Conceptualization, formal analysis, funding acquisition, and writing – original draft preparation; **SDH:** Data curation, formal analysis, software, and writing – review and editing; **SMB:** Formal analysis, investigation, and writing – original draft preparation; **NP:** Conceptualization and investigation; **DGW:** Conceptualization, funding acquisition, resources, supervision, and writing – original draft preparation.

## Acknowledgements

Thank you to the Vanderbilt Mouse Neurobehavior Lab for assistance with tube test assays and equipment. MAD was supported by a K99 from NIAAA (AA031509). CME was supported by an F32 from NIAAA (AA031404). NP was supported by an F30 from NIAAA (AA029599), a T32 from NIGMS (GM007347), and an R01 Diversity Supplement from NINDS (NS102306). The research was supported by an R37 (DGW, AA019455) and P60 (DGW, AA031124) from NIAAA.

---

**Supplemental Figure 1.**
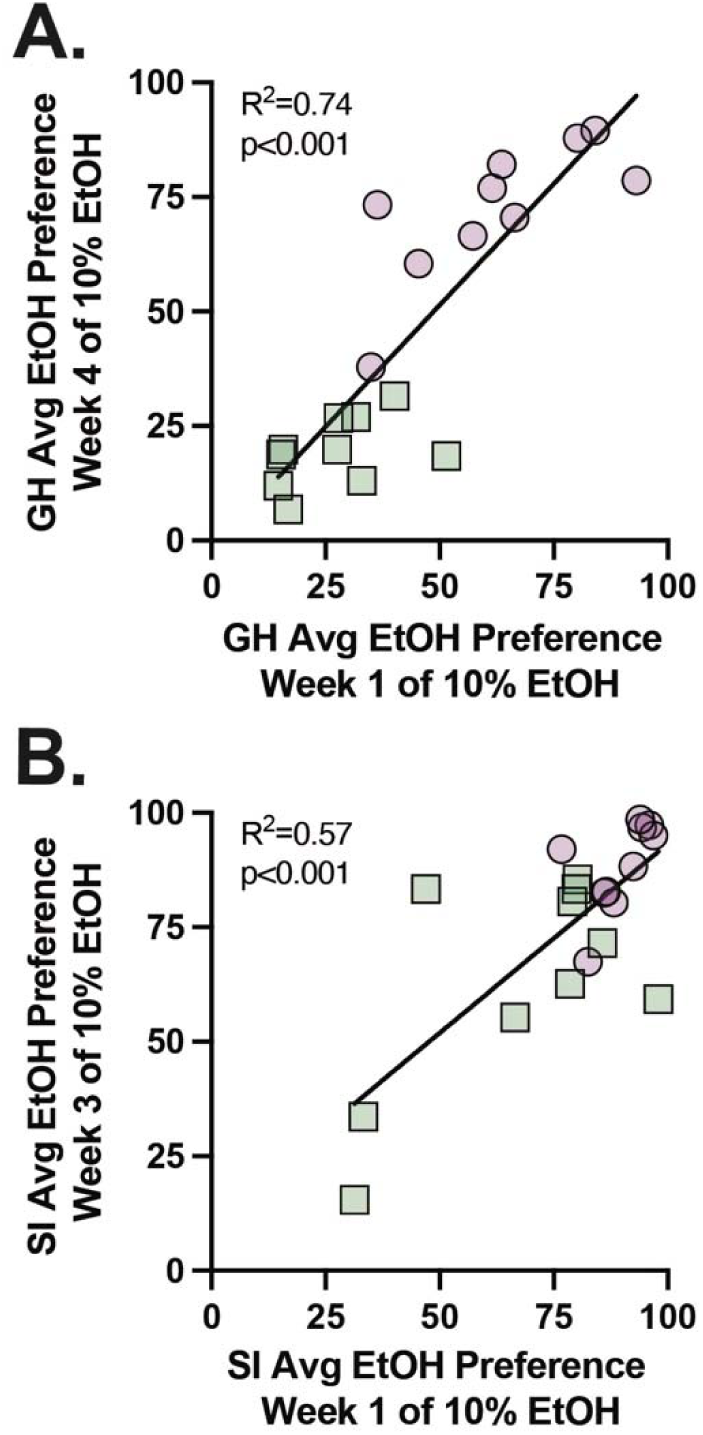
Drinking preference reflected stable individual drinking patterns. **A.** A significant positive correlation was observed between the first and last week of 10% ethanol access in group housed mice (n=20, simple linear regression). **B.** Socially isolated mice exhibited a significant positive correlation between ethanol preference measured during the first and last weeks of 10% ethanol access. (n=20, simple linear regression).

**Supplemental Figure 2.**
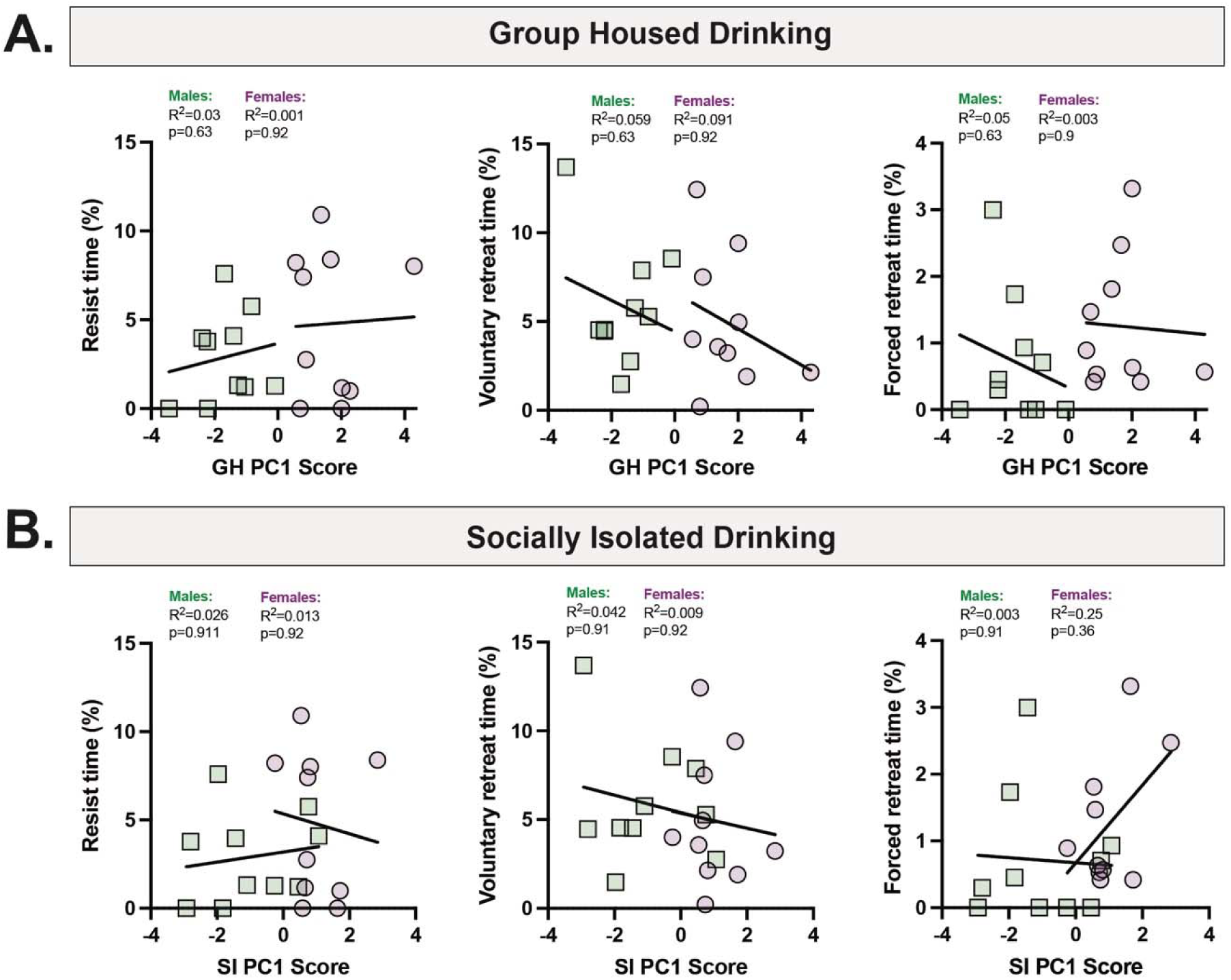
Defensive tube test metrics were not associated with PCA-defined drinking phenotypes in either group-housed or socially isolated drinking environments. **A.** Defensive tube test metrics of percent time resisting, voluntarily retreating, and forced retreating were not significantly correlated with principal component scores during group housed drinking in male or female mice (n=10/sex, Simple linear regression with FDR correction). **B.** In both male and female mice, defensive tube test were not significantly associated with principal component scores from socially isolated drinking behavior (n=10/sex, Simple linear regression with FDR correction).

**Supplemental Figure 3.**
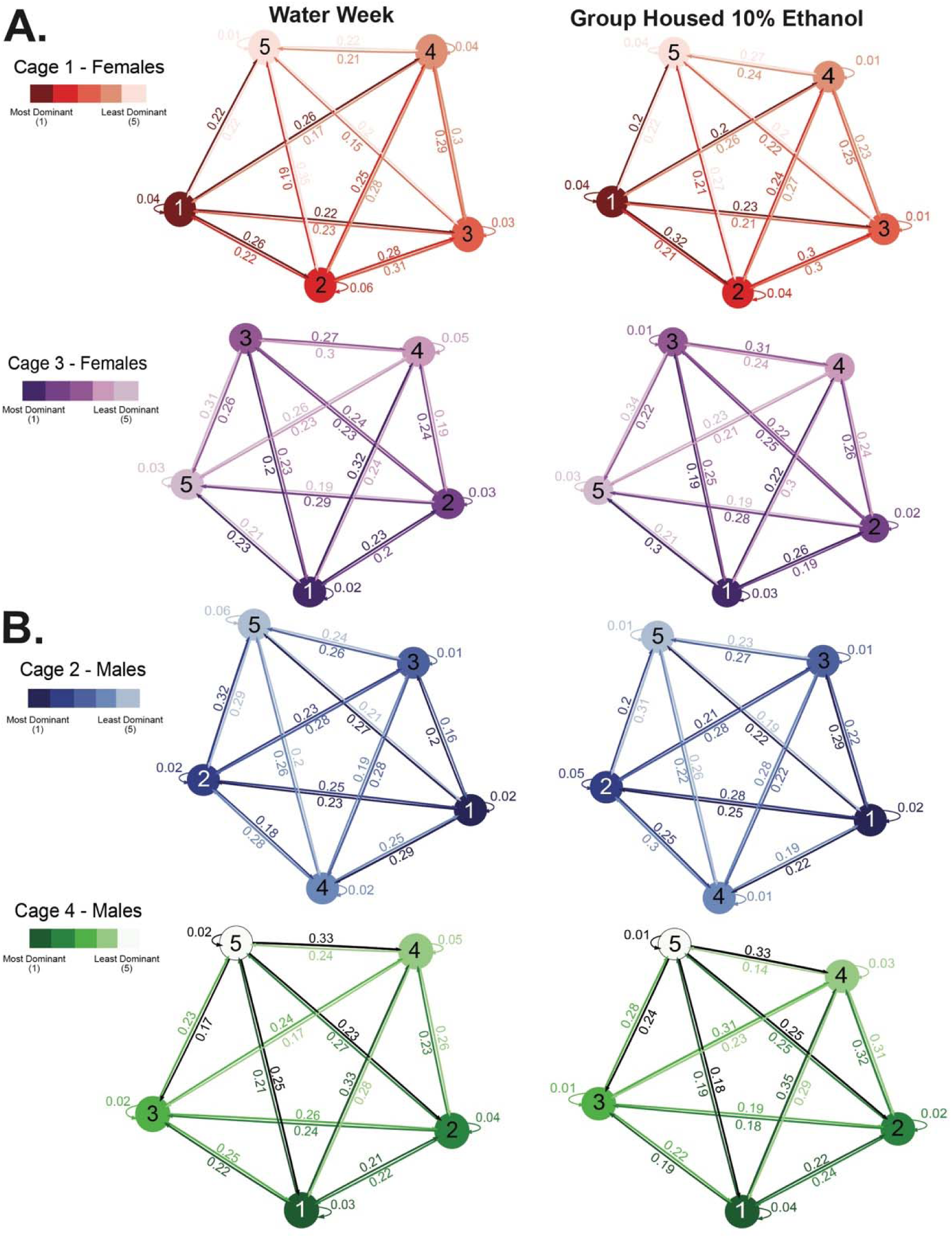
Drinking order as assessed by meal pattern analysis was not predicted by social rank. **A.** Markov modeling of meal drinking order in female mice did not display a relationship between social rank and drinking order. **Left:** Water week; **Right:** Last week of group housed 10% ethanol access (n=5 mice/cage; n=2 cages/sex). **B.** In male mice, Markov analysis of meal drinking order showed no association between social rank and drinking order. **Left:** Water week; **Right:** Last week of group housed 10% ethanol access (n=5 mice/cage; n=2 cages/sex)

